# *In vivo* proximity ligation reveals endogenous candidate interactors of Neurexin’s intracellular domain

**DOI:** 10.1101/2022.09.27.509791

**Authors:** Marcos Schaan Profes, Araven Tiroumalechetty, Neel Patel, Stephanie S. Lauar, Simone Sidoli, Peri T. Kurshan

## Abstract

Neurexins are highly-spliced transmembrane cell adhesion molecules that bind an array of partners via their extracellular domains. However, much less is known about the signaling pathways downstream of neurexin’s largely-invariant intracellular domain. C. elegans contains a single neurexin gene that we have previously shown is required for presynaptic assembly and stabilization. To gain insight into the signaling pathways mediating neurexin’s presynaptic functions, we employed a proximity ligation method, endogenously tagging neurexin’s intracellular domain with the promiscuous biotin ligase TurboID, allowing us to isolate adjacent biotinylated proteins by streptavidin pull-down and mass spectrometry. We compared our experimental strain to a control strain in which neurexin, endogenously tagged with TurboID, was dispersed from presynaptic active zones by the deletion of its C-terminal PDZ-binding motif. Using this approach we identified both known and novel intracellular interactors of neurexin, including active zone scaffolds, actin-binding proteins (including almost every member of the Arp2/3 complex), signaling molecules, and mediators of RNA trafficking, protein synthesis and degradation, among others. Characterization of mutants for candidate neurexin interactors revealed that they recapitulate aspects of the *nrx-1* mutant phenotype, suggesting they may be involved in neurexin signaling. Finally, to investigate a possible role for neurexin in local actin assembly, we endogenously tagged its intracellular domain with actin depolymerizing and sequestering peptides (DeActs), and found that this led to defects in active zone assembly.

## Introduction

The proper formation of synaptic connections underlies our brain’s ability to form appropriate neuronal circuits, and defects in this process lead to neurodevelopmental and neuropsychiatric disorders. Synaptic cell-adhesion molecules (sCAMS) are thought to play a role in both the specificity of this process, by selecting appropriate synaptic partners^1–3^, and in the stabilization and functional maturation of nascent synapses^4,5^.

Neurexins constitute a family of presynaptic CAMs that are highly associated with autism and schizophrenia^6^, and are thought to function as central “hubs” of trans-synaptic interaction^7^. The synaptogenic activity of neurexin was initially demonstrated by showing that binding to its canonical binding partner neuroligin could induce the formation of hemi-presynapses in cultured neurons^8–10^. The human genome encodes three neurexin genes, which together can be expressed as ∽4,000 different splice isoforms^11,12^. These isoforms contain a mostly invariant intracellular domain (ICD) responsible for a largely uncharacterized downstream intracellular signaling pathway: the intracellular C-terminal PDZ-binding motif (PBM) of neurexin interacts with the synaptic vesicle protein synaptotagmin as well as the scaffolding proteins Cask and Mint^13–17^. In addition, *Drosophila* neurexin has been shown to interact with the active zone protein SYD-1^18^ as well as the actin binding protein spinophilin^19^.

*C. elegans* contains a single neurexin gene (nrx-1) that encodes both long and short isoforms^20,21^. The long isoforms of NRX-1 have been implicated in neurite outgrowth, synapse specificity and postsynaptic organization^22,23^, while the short isoform is sufficient for presynaptic maturation and stability^21^. Using markers for presynaptic assembly including the synaptic vesicle (SV)-associated protein RAB-3 and the active zone (AZ) protein clarinet (CLA-1; homolog of vertebrate AZ protein Piccolo^24^), we have previously shown that *C. elegans* NRX-1 stabilizes nascent synapses and is required for their morphological and functional maturation^21^. However, the downstream signaling pathways responsible for these functions remain unknown.

To better understand the molecules that might mediate neurexin’s presynaptic role in synapse stabilization and maturation we have employed the enzyme-catalyzed proximity-labeling approach TurboID^25^. This method utilizes the promiscuous biotin ligase BirA, fused to a protein of interest, to allow for biotinylation of target proteins within a radius of a few nanometers. Biotinylated proteins are pulled down with streptavidin and identified by mass spectrometry. Unlike traditional biochemical approaches, this method does not require interacting proteins to remain in complex during purification, a particular advantage when studying transmembrane proteins or looking for transient interactions.

To identify proteins that interact with neurexin intracellularly we used CRISPR gene editing to endogenously tag the neurexin intracellular domain with TurboID, and confirmed that this does not affect neurexin function *in vivo*. Streptavidin pull-downs and mass spectrometry were used to identify biotinylated proteins. We then compared our results to three different negative controls: a wild type strain lacking any TurboID protein, a strain over-expressing cytosolic TurboID pan-neuronally, and a strain in which TurboID was endogenously tagged to NRX-1, but in which the PBM of NRX-1 had been deleted leading to a de-clustering of NRX-1 from presynaptic active zones. We conclude that the latter strain is the most appropriate negative control, the former two being too permissive or too restrictive, respectively. Using this control, we have generated a list of potential NRX-1 interactors, including both known and novel binding partners. These include presynaptic active zone proteins as well as many proteins involved in remodeling of the actin cytoskeleton. We characterized mutants for a subset of these proteins, and discovered that they recapitulate aspects of the *nrx-1*mutant phenotype, suggesting they may be involved in neurexin signaling. Finally, to directly assess the role of actin polymerization in neurexin’s presynaptic function, we fused bacterially-derived actin-depolymerizing or sequestering peptides to neurexin’s intracellular domain and found that this resulted in a pronounced reduction in active zone size.

## Results

### Endogenous tagging and validation of neurexin with intracellular TurboID

Neurexin mutants have a defect in presynaptic stability and thus are more susceptible to extrinsic inhibitory cues, the result of which is that they have fewer active zones clusters, particularly at the edges of the synaptic domain (where inhibitory cues are highest)^21^. They also exhibit an increase in the number of small, highly-mobile synaptic vesicle precursor packets in the asynaptic region of the axon^21^. These dual phenotypes allow us to assess neurexin function using a transgenic marker that expresses both a fluorescently tagged active zone protein (Clarinet, or CLA-1^24^) and a synaptic vesicle protein (RAB-3) in the DA9 motor neuron in the tail of the worm^21^.

The ICD of neurexin is largely uncharacterized and contains few sequence motifs, with the notable exception of a C-terminal PDZ-binding motif (PBM). To identify an appropriate location within neurexin’s ICD in which to insert the TurboID biotinylating enzyme (BirA), we considered three options: (1) just after the transmembrane domain, (2) just before the PBM, and (3) at the very C-terminus with an extra-long linker (Fig. 1A). We generated rescue constructs of each and assayed their ability to rescue the neurexin null phenotype, using the marker described above. Insertions at the first two locations were able to rescue the null phenotype (data not shown), however the third (C-terminal) option failed in rescuing the phenotype, thus it was discarded. We proceeded to generate TurboID endogenous CRISPR knock-in strains of the endogenous neurexin locus at both other two ICD locations (see materials and methods). In contrast to our over-expression rescue experiments, the first (post-transmembrane domain) led to a neurexin null phenotype (data not shown), indicating that the endogenous insertion had abrogated neurexin’s function. However, the second (pre-PBM; Fig. 1A,B), resulted in wild type presynaptic development (Fig. 1C-E, suggesting that the insertion of TurboID at this location did not impact neurexin function in presynaptic assembly and stability.

**Figure 1.**
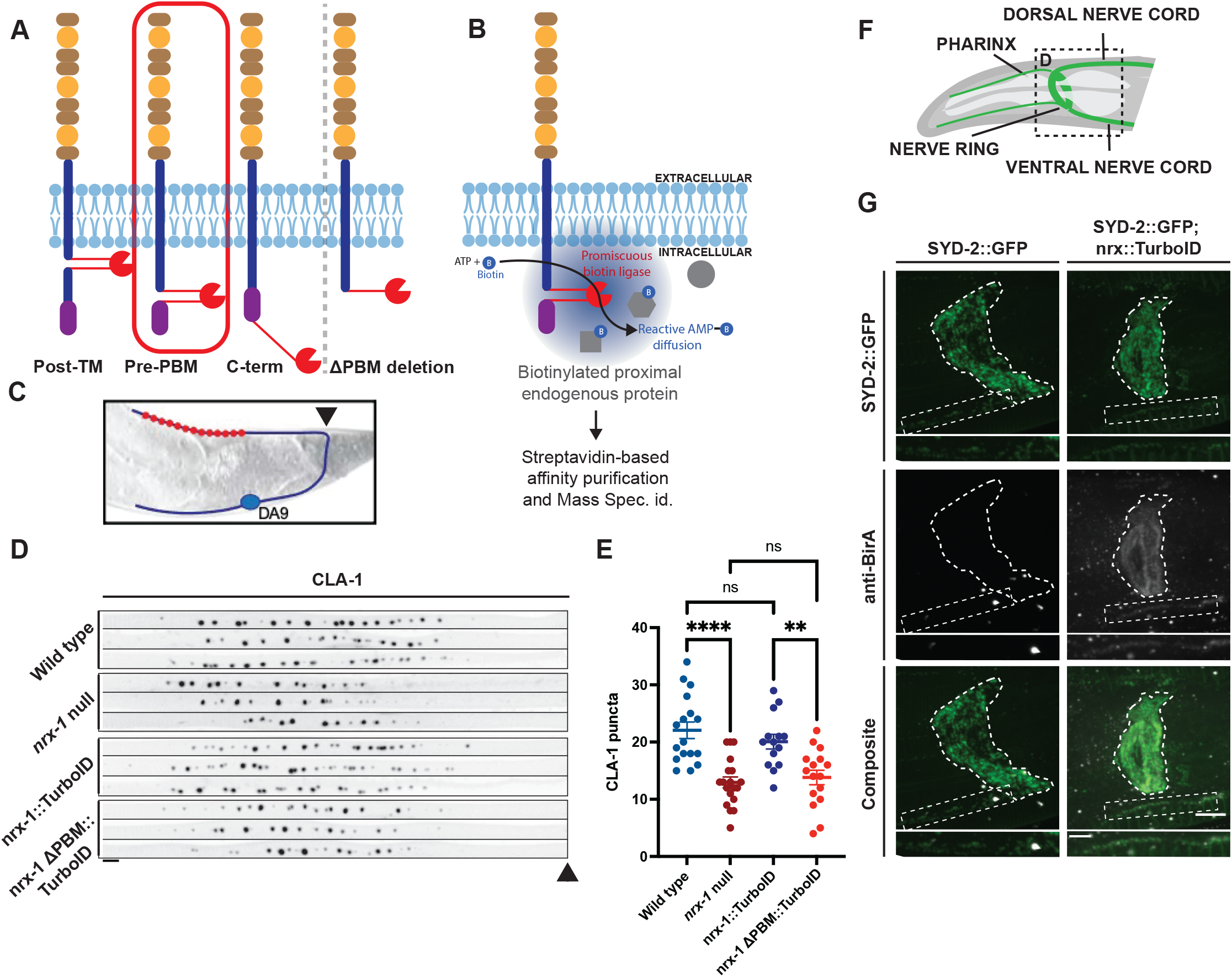
Generation and validation of an endogenously-tagged neurexin-TurboID strain and control. **A**. Left: Schematic depicting several insertion sites of TurboID enzyme that were assessed, with the final chosen and validated site circled in red. Right: Schematic of the neurexin-ΔPBM-TurboID control strain. **B**. Schematic of the rationale and workflow for the proteomics screen. **C**. Schematic of the DA9 motor neuron used to assess presynaptic assembly phenotypes. Arrowhead points to where cropped images begin in D. **D**. Straightened images of CLA-1-GFP puncta in the DA9 synaptic domain across different genotypes. Scale bar: 4μm. **E**. Quantification of CLA-1 puncta number in the indicated genotypes reveals that our experimental strain (neurexin-TurboID) does not impact neurexin function, but our ΔPBM negative control strain does. **F**. Schematic of the synapse-rich nerve ring in the head of the worm. **G**. Immunohistochemistry using anti-BirA antibody compared to GFP fluorescence of endogenously-tagged active zone protein SYD-2 in the nerve ring and nerve cord (insets) reveals synaptic localization of our endogenously-tagged neurexin-TurboID. Scale bars: 10μm for nerve ring images and 5μm for insets.

We further validated this strain by performing immunocytochemistry on our TurboID-tagged neurexin strain, using antibodies against BirA, and comparing the pattern of expression to another endogenously-tagged presynaptic active zone protein, SYD-2/Liprin-α^26^. Expression of both neurexin-TurboID and SYD-2-GFP colocalized well in the synapse-rich region of the nerve ring (Fig. 1F,G), as well as in the individual puncta of the nerve cord (Fig. 1G, insets), indicating that neurexin-TurboID was localizing appropriately to presynaptic active zones.

Previous TurboID experiments in *C. elegans* have made use of a negative control strain in which cytosolic BirA is over-expressed in the tissue of interest through the use of an integrated multi-copy array^27^. To generate a more appropriate and highly specific negative control strain for our TurboID proteomics experiments, we genetically removed the PBM from our endogenously-tagged neurexin-TurboID strain (see methods and Fig. 1A), as this leads to the de-clustering of neurexin and its dispersal along the cell surface (our unpublished results). Indeed, the deletion of the PBM in the neurexin-TurboID strain led to a synaptic assembly phenotype similar to that of the neurexin null mutant (Fig. 1D,E), indicating that neurexin’s localization at active zones is critical to its function in presynaptic assembly and stability.

### Proteomics results and comparison to multiple negative control strains

To identify candidate proteins that may interact with neurexin’s intracellular domain, we set out to perform proteomics analysis of our endogenous neurexin-TurboID strain, compared to three different negative control strains: wild type (N2), which contains no BirA enzyme; the pan-neuronally over-expressed cytosolic TurboID strain (wyIs687); and our newly generated neurexin-ΔPBM-TurboID strain (Fig. 2A-D). Developmentally synchronized worms enriched for adults were grown on standard bacterial medium (OP50, which contains low levels of biotin). Two hours prior to their lysis, half of the replicates of each strain were incubated on media supplemented with 1mM of biotin.

**Figure 2.**
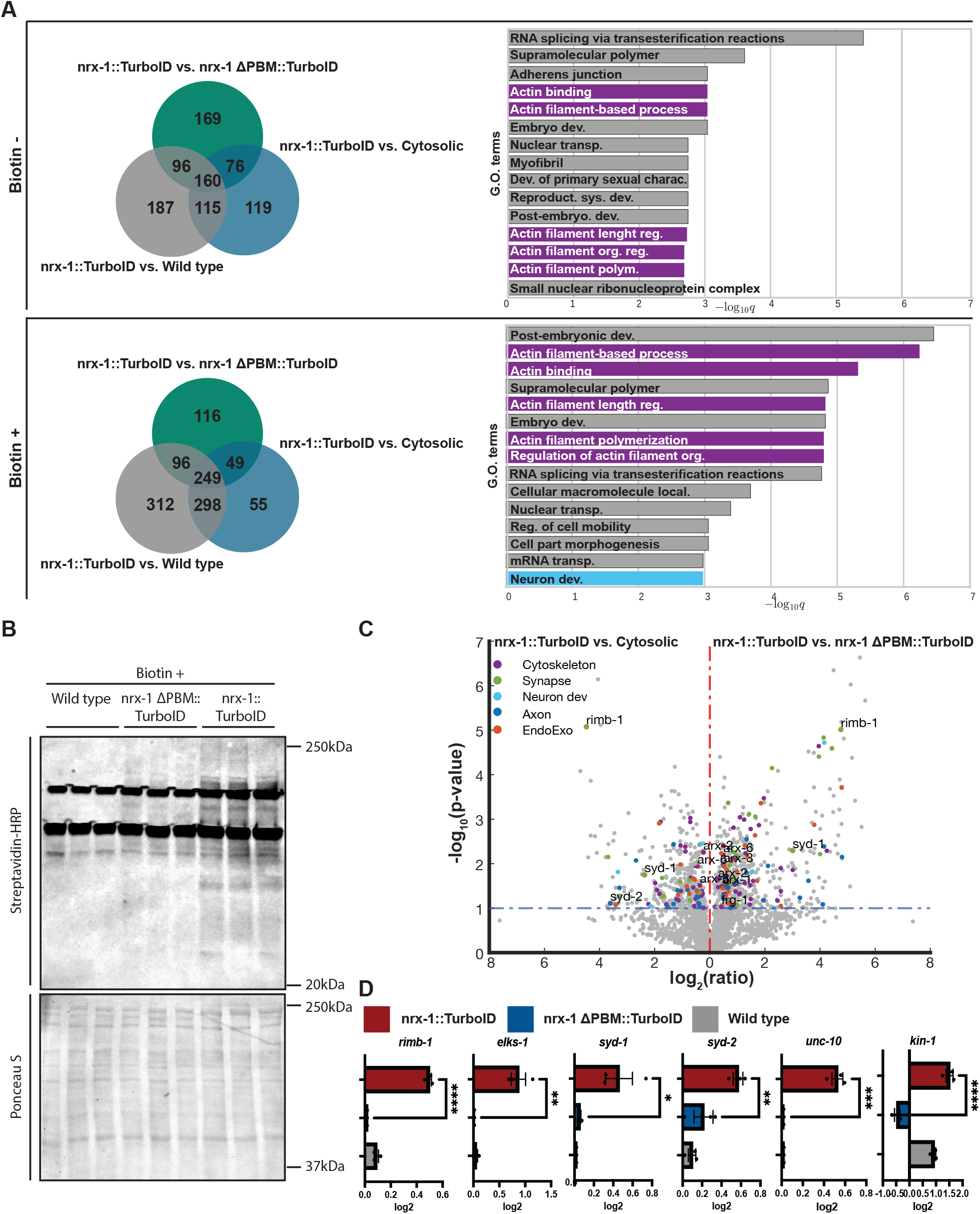
Comparison of proteomics results between multiple negative control strains. **A**. Left: Venn diagrams showing proteins enriched in our experimental strain (neurexin-TurboID) compared to three different negative control strains (wild type N2, pan-neuronal cytosolic TurboID, and neurexin-ΔPBM-TurboID), in both basal and enriched Biotin conditions. Right: GO terms of most highly enriched genes in comparison to neurexin-ΔPBM-TurboID in both basal and enriched Biotin conditions. **B**. Western blot of biotinylated proteins in our experimental strain (neurexin-TurboID, right columns) compared to two controls (wild type, left columns and neurexin-ΔPBM-TurboID, middle columns) in the enriched Biotin condition as probed by streptavidin-HRP. Total protein levels (as assessed by Ponceau staining, lower blot) were used as a loading control. **C**. Volcano plot of genes corresponding to the proteins enriched in our experimental strain (neurexin-TurboID) compared to two negative controls (over-expressed cytosolic pan-neuronal TurboID on the left and neurexin-ΔPBM-TurboID on the right). **D**. Absolute enrichment values transformed in log_2_ compared to two controls (wild type and neurexin-ΔPBM-TurboID) for known active zone components likely to be closely physically associated with neurexin’s intracellular domain.

The lysates from each strain/condition were then used to perform streptavidin pull-downs to isolate biotinylated proteins (see methods and Fig. 1B). Following pull-downs, we performed Western blots to assess and validate our purification and to control for BirA protein biotinylation (Fig. 2B and Supplementary Fig. 1A). Total protein levels (as assessed by Ponceau staining; Fig. 2B and Supplemental Fig. 1A) were used as a loading control and biotinylated proteins were assessed by immunoblotting with streptavidin-HRP. The experimental strain, neurexin-TurboID, showed increased biotinylated protein levels and resulted in more easily identified specific bands following streptavidin immunoblotting (Fig. 2B) when compared to the controls. This was particularly noticeable in the added-biotin conditions (compare Fig. 2B to Supplemental Fig. 1A), suggesting an increase in specificity in this condition. Moreover, when comparing the biotin-enriched condition to the basal condition, we saw an increase in the number of candidate genes with gene ontology (GO) terms predicted to be relevant to neurexin function (e.g. synapse, neuron development, axon, endo/exocytic-related, and cytoskeleton; Supplemental Fig 1B). Additionally, these hits displayed higher fold-change and/or p-value compared to the non-biotin enriched samples, again suggesting increased specificity in the biotin-enriched condition.

Following streptavidin pull-downs, samples were submitted for mass spectrometry at the Einstein proteomics core facility. The dataset was then processed with logarithm transformation (to fit the data to a normal distribution, as proteomics data have a positively skewed distribution), normalized to total protein levels, and missing values were imputed (replaced using the Probabilistic Minimum Imputation for label-free data, as described in ^28^).

We compared the proteomics results of our experimental strain (neurexin-TurboID) to our three negative control strains (wild type, cytosolic TurboID, and neurexin-ΔPBM-TurboID), in both biotin conditions, and constructed venn diagrams of the overlapping enriched hits in the samples, using a 90% confidence threshold in t-tests for including candidates (Fig. 2A). Using a combination of Gene Ontology (GO) (Fig. 2A) and volcano plots (Fig. 2C) to compare these enriched hits in our experimental strain relative to either the overexpressed cytosolic TurboID or our neurexin-ΔPBM-TurboID controls, we found a greater number and enrichment of relevant neuronal, synaptic and cytoskeletal terms in the latter condition. This was particularly true for the biotin-enriched samples, where these hits were both further enriched and higher up on the gene ontology list (Fig. 2A). We interpreted this as indicating that the over-expressed cytosolic TurboID, due to its high expression level, may obscure real neurexin interactors. This might especially be the case for interactors that are themselves highly expressed throughout the cell, such as cytoskeletal proteins, thus making this strain too stringent a negative control. For example, GO analysis of candidate interactors obtained using the neurexin-ΔPBM-TurboID as a control revealed an increase in actin-related terms as compared with using the cytosolic TurboID control (Fig. 2C). Identification of several components of the presynaptic active zone, including RIMB-1, ELKS-1, SYD-1, SYD-2/Liprin-α and UNC-10/Rim (Fig. 2D), gave us confidence in the specificity of our results. In addition, we found enrichment of the *C. elegans* PKA ortholog KIN-1 (Fig. 2D). The mammalian version of this protein has been implicated in regulating presynaptic potentiation downstream of neurexin^29^. Overall, we concluded that our specific endogenous control strain (neurexin-ΔPBM-TurboID) is the most appropriate control strain, since it is expressed off of the endogenous neurexin promoter (and therefore likely at similar levels to our experimental strain) and differs only in its subcellular localization pattern (loss of synaptic enrichment), and we proceeded in our analysis using that comparison.

### Neurexin interactions with novel proteins and signaling pathways

Having determined the most appropriate negative control, we began our analysis of candidate interacting proteins revealed by the proteomics analysis. To select those, we again used a 90% confidence threshold in a t-test for including candidates. We found candidate interactors that fell into several broad classes: active zone proteins (Fig. 2D), cytoskeletal-associated proteins, in particular actin-related proteins, including almost all the members of the actin-nucleating Arp2/3 complex (arx genes in *C. elegans*), other actin-associated proteins (frm-1, frm-4, frg-1,hum-4, ctn-1, pfn-1, dbn-1, pkc-3, twf-2, unc-115, and unc-60), additional synaptic proteins (ric-4, ddi-1, sax-7), as well as those involved in autophagy and other categories (Fig. 3 and data not shown). We plotted the absolute values transformed in log_2_ for each protein in each condition (experimental, ΔPBM, and wild type strains) for easier comparison (Fig. 3).

**Figure 3.**
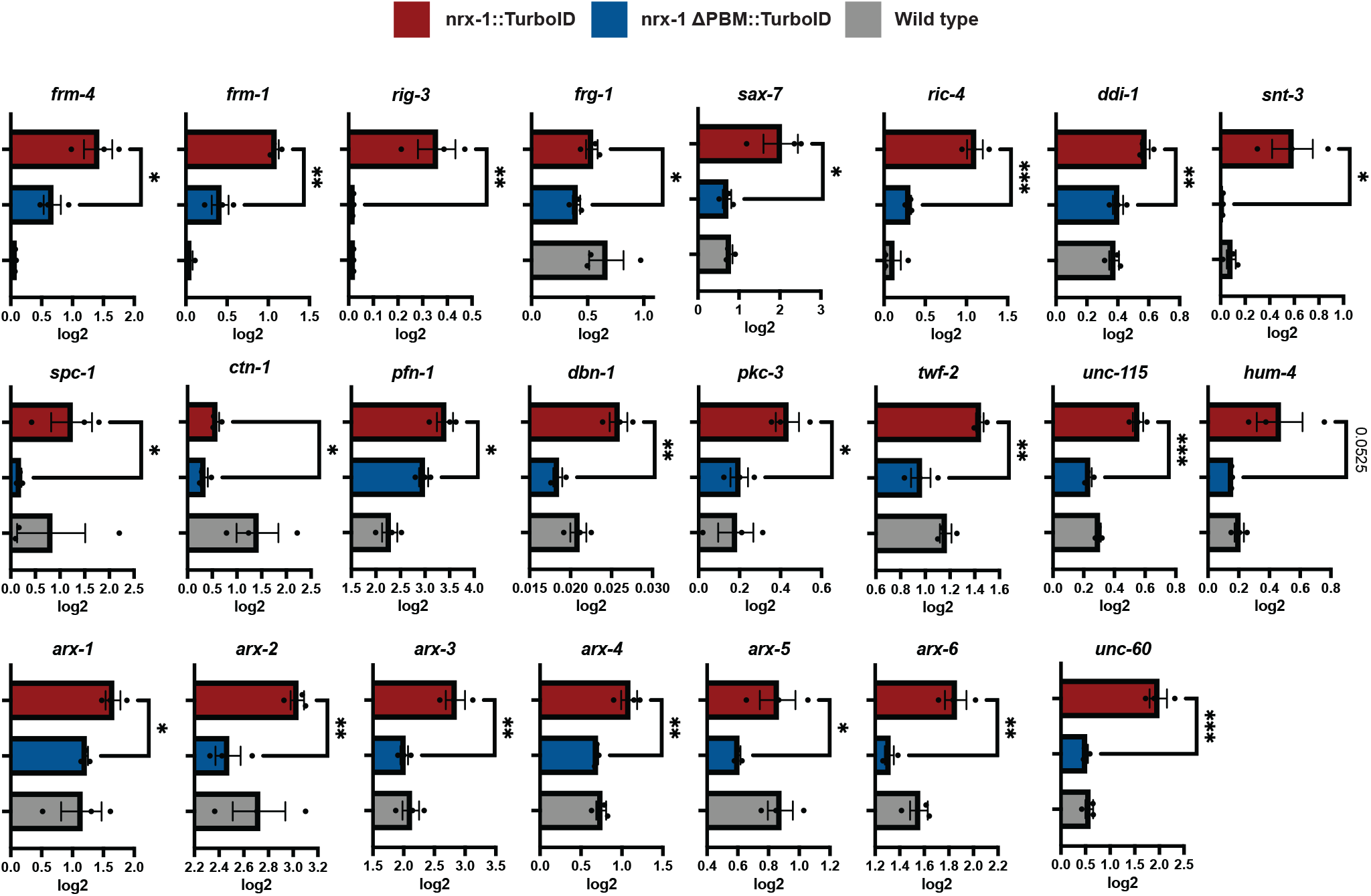
Candidate neurexin interactors in multiple molecular pathways. Absolute enrichment values transformed in log_2_ compared to two controls (wild type and neurexin-ΔPBM-TurboID) for a subset of genes of interest.

### Mutants of candidates from proteomics screen partially phenocopy neurexin mutants and have varied effects on synapse assembly/stability

Mutants for several genes identified in our screen (Fig. 4A-C) were obtained from stock centers (see strain list in materials and methods), crossed to our synaptic marker strain and assessed for presynaptic assembly defects. These include *frm-4, frg-1, ctn-1, hum-4, sma-1, and rig-3*. Frm-4, encodes a FERM domain-containing protein predicted to be involved in actomyosin structure organization. Frg-1 (ortholog to FSH muscular dystrophy Region Gene 1), ctn-1 (alpha-CaTuliN), hum-4 (heavy chain of an unconventional myosin) and sma-1 (SMAll) all encode proteins that are predicted to enable actin filament binding activity. Rig-3 (neuRonal IGCAM) encodes an adhesion molecule located in axons and synapses.

**Figure 4.**
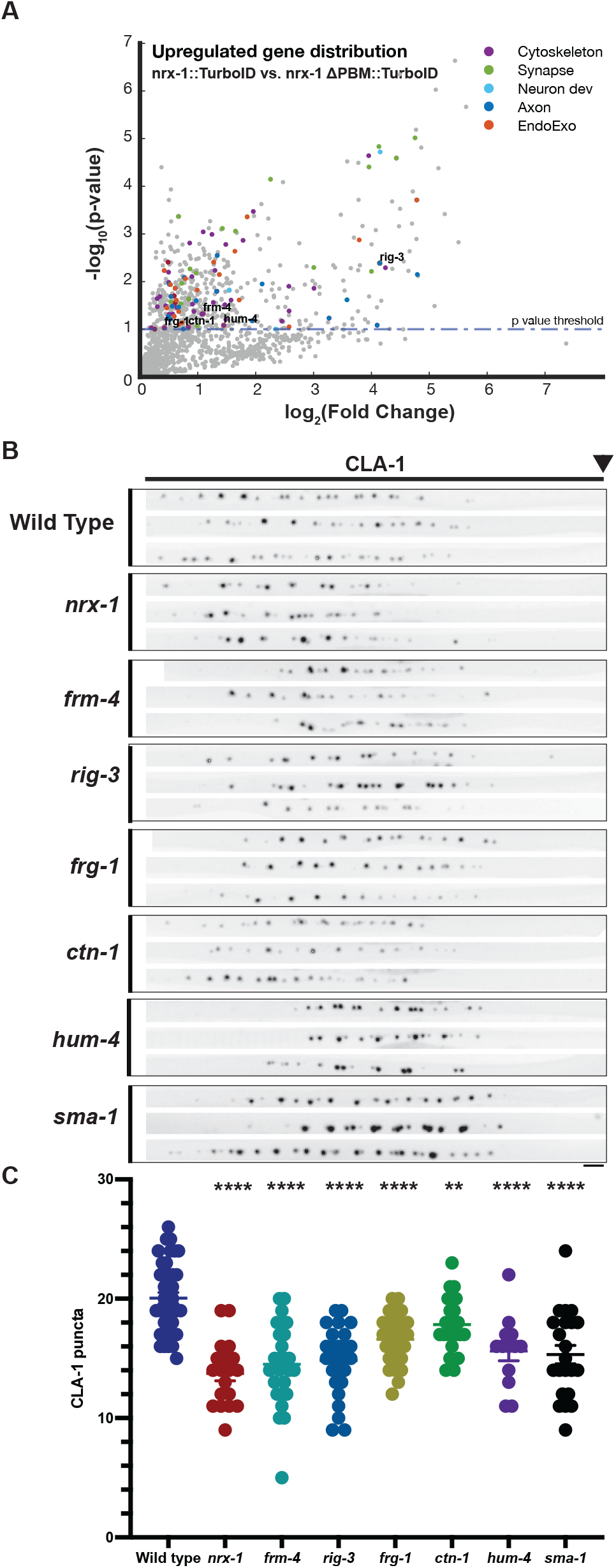
Mutants of candidate neurexin interactors partially phenocopy neurexin mutant phenotypes. **A**. Semi-Volcano plot of genes corresponding to the proteins enriched in our experimental strain (neurexin-TurboID) compared to control (neurexin-DPBM-TurboID), replotted from Fig. 2C, but with selected candidate interactor genes highlighted to show their relative enrichment within the dataset. **B**. Straightened images of CLA-1-GFP puncta in the DA9 synaptic domain across different genotypes. Scale bar: 4μm. **C**. Quantification of CLA-1 puncta number in the indicated genotypes.

Compared to wild type animals, *nrx-1* null mutants exhibit a ∽30% reduction in the number of active zones (CLA-1 puncta), primarily within the proximal synaptic domain, as well as an increase in small, asynaptic vesicle precursors (RAB-3 puncta) in the axon commissure (Fig. 4B,C, Supplementary Fig. 2B, and ^21^). Notably, several of the mutants of candidate interactors exhibited similar defects in presynaptic assembly and stability, including a reduction in the number of synaptic CLA-1 puncta (Fig. 4B,C), as well as an increase in asynaptic RAB-3 puncta (Supplementary Fig. 2B). *Frm-4, rig-3, frg-1* and *hum-4* mutants showed a pronounced reduction in CLA-1 puncta in comparison to wild type. Interestingly, this reduction was mostly seen in the distal portion of the axon. On the other hand, *ctn-1* showed a more modest reduction, but this was seen mostly in the proximal region of the axon, similar to the region most affected by loss of neurexin (Fig. 4B and C). *Sma-1* mutants, although also displaying a significant reduction in CLA-1 puncta, were much smaller in size, complicating our analysis. Interestingly, only *frm-4* and *hum-4* mutants also recapitulated the increase in asynaptic RAB-3 seen in the *nrx-1* null strain (Supplemental Fig. 2B). Altogether, this data suggests that NRX-1 may function upstream of several different pathways controlling presynaptic assembly and stability.

### Neurexin’s intracellular domain may regulate presynaptic actin organization and/or polymerization

Our gene ontology analysis showed a prominent enrichment in actin-related proteins including almost all the members of the actin-nucleating Arp2/3 complex (arx genes in *C. elegans*) and other actin-associated proteins (frm-1, frm-4, frg-1,hum-4, ctn-1, pfn-1, dbn-1, pkc-3, twf-2, unc-115, and unc-60; Fig. 2, 3 and Supplementary Figure 3). Due to the importance of the actin cytoskeleton in presynaptic structure and organization, redundant signaling pathways are likely involved, making single-mutant analysis hard to interpret. Moreover, many actin proteins are essential in worms and their mutants therefore lethal. To understand whether neurexin may mediate very local changes in actin organization, we decided to employ a strategy aimed at specifically perturbing actin polymerization surrounding neurexin’s intracellular domain. DeActs are a class of bacterially-derived, genetically encoded actin-modifying polypeptides, that can induce actin disassembly in eukaryotic cells^30^. Using CRISPR/Cas9, we endogenously tagged neurexin’s ICD with the DeAct Gelsolin segment 1 (GS1), a ∽120-amino-acid domain that sequesters actin monomers, placing it in the same location that we had previously inserted TurboID (Fig. 5A). We found that in nrx-1::GS1, the number of active zones marked by the active zone scaffold Clarinet (CLA-1) was unaltered, however the average size of CLA-1 puncta was decreased (Fig. 5B-C), a defect in in active zone assembly even more pronounced than that found in neurexin mutants. Altogether, our data suggest that neurexin may mediate presynaptic assembly in part by interacting with factors regulating actin polymerization and/or organization.

**Figure 5.**
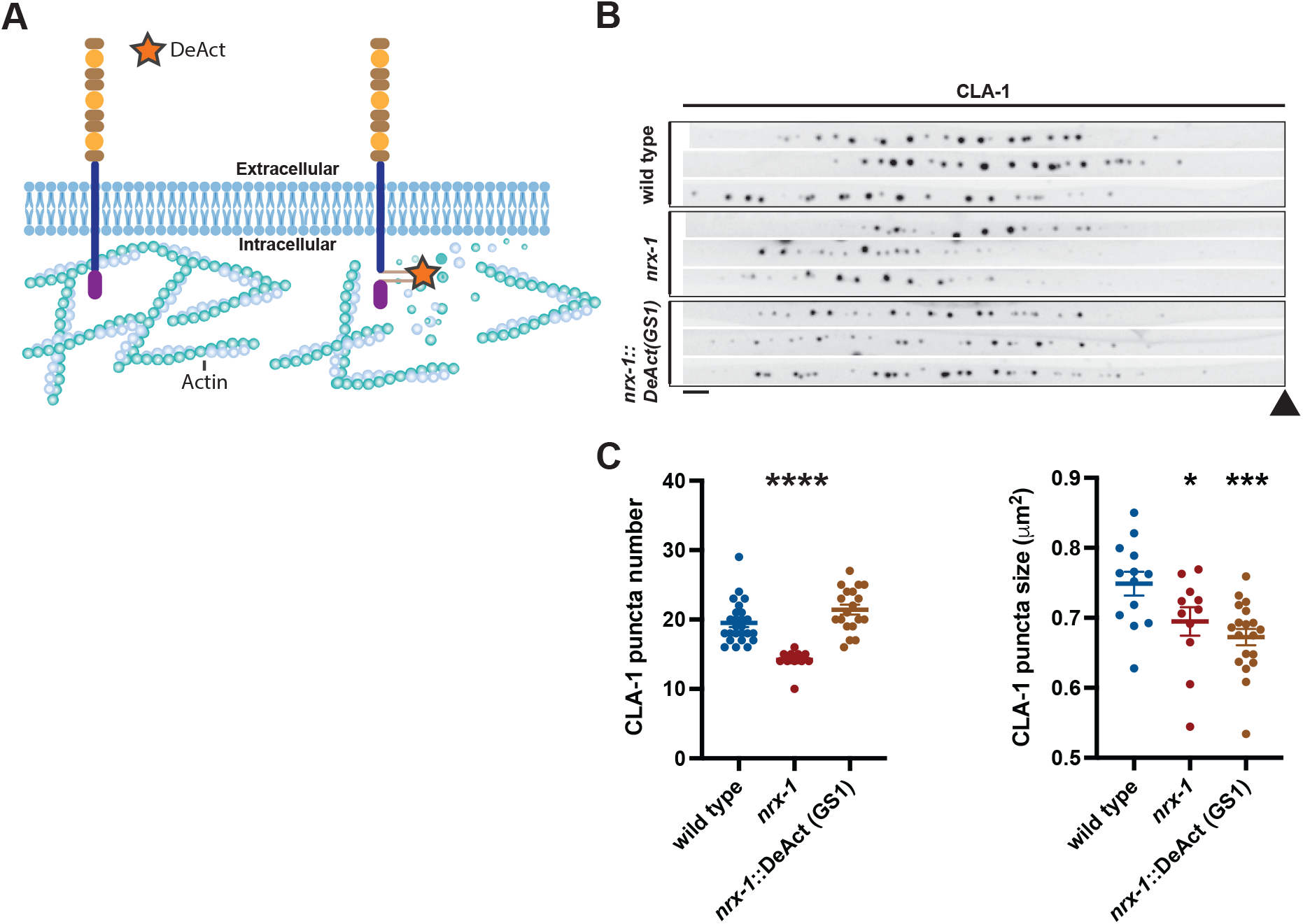
Endogenously-tagging neurexin’s ICD with the actin depolymerizing peptide DeAct (GS1) leads to a reduction in active zone size. **A**. Schematic depicting the insertion site of DeAct tool Gelsolin segment 1 (GS1). **B**. Straightened images of CLA-1-GFP puncta in the DA9 synaptic domain across wild type, *nrx-1 null* and *nrx-1::*DeAct(GS1) genotypes. Scale bar: 4μm. **C**. Quantification of CLA-1 puncta number and size in the indicated genotypes.

## Discussion

This is, to our knowledge, the first proximity labeling experiment conducted to identify interactors of neurexin’s intracellular domain, a region common to all neurexin genes and isoforms and thus critical for mediating neurexin signaling in all neuronal contexts. Moreover, we have conducted this analysis using endogenously-tagged neurexin *in vivo*, thus retaining the appropriate cellular context and abrogating any effects of over-expression. Careful selection and validation of the endogenous insertion site resulted in generation of an experimental strain with wild type neurexin function, while an analysis of several possible negative control strains led to the selection of the most appropriate one. We have identified both known and novel candidate interactors of neurexin’s intracellular domain, revealing unknown roles for these proteins in presynaptic assembly and stability. In particular, we have identified a likely role for neurexin in actin nucleation, due to the identification of almost every member of the actin-nucleating Arp2/3 complex in our proteomic screen results.

Arp2/3 is crucial for regulation of both the initiation of actin polymerization and organization of the resulting filaments into branched networks^31^. Actin polymerization has been shown to be required for the development of synaptic structures and the clustering of synaptic vesicles within presynaptic boutons^32^. In *Drosophila*, neurexin has been shown to interact genetically with the actin binding protein spinophilin^19^. However, a direct connection between neurexin signaling and actin polymerization has not yet been reported. Although more studies are required to validate the link between neurexin and actin polymerization, the enrichment of actin-binding and actin-nucleating proteins in our proteomics results coupled with the pronounced effect on active zone size obtained by fusing the DeAct peptide GS1 to neurexin’s intracellular domain suggests that neurexin may play a role in actin modification at the active zone. Taken together, our data suggest an important link between neurexin and presynaptic actin organization to mediate presynaptic assembly, stability and function.

Several of the synaptic proteins enriched in our proteomic analysis have not been previously linked to neurexin signaling. Ric-4, which is involved in cholinergic synaptic transmission in *C. eleganssie*^33^, is an ortholog of human SNAP25, which have been associated with Down syndrome^34^. Ddi-1 has been implicated in negative regulation of synaptic assembly in *C. elegans*, with its mutants displaying a significant increase in synaptic density along the dorsal nerve cord^35^. The immunoglobulin cell adhesion molecule sax-7 has been implicated in maintaining placement of neurons and their axons^36^, and more recently genetically linked to RAB-3, suggesting a possible function in synaptic vesicles exocytosis^37^. None of the mutants we analyzed perfectly recapitulated the *nrx-1* null mutant phenotype, suggesting that neurexin may function as a signaling hub upstream of several different signaling pathways for synapse assembly, stability and maturation. Altogether, our data suggests that neurexin may interact with several important structural, organizational and functional synaptic players to mediate presynaptic development through distinct signaling pathways.

Interestingly, we also found hits in other classes of proteins, including those involved in the direct regulation of exocytosis (including SNARE proteins), in autophagy, in calcium signaling, as well as various kinases and axon guidance molecules. This suggests that there may be non-canonical functions of neurexin that together characterize its complex role in presynaptic regulation.

An important contribution of this study is our in-depth analysis of several different conditions and negative control strains. In order to be useful, proteomic screens must have a good signal-to-noise ratio. Our goal in comparing our experimental strain to three different negative control strains, including one generated specifically for this experiment, was to identify the comparison with the best ratio. We concluded that comparison to a wild type strain (no TurboID) was too permissive, while comparison to an over-expressed cytosolic TurboID was too restrictive. Generation of a specific control strain in which TurboID was still tethered to neurexin and expressed at endogenous levels off the endogenous promoter, but in which neurexin’s clustering at the active zone was specifically abrogated, furnished us with the greatest enrichment of expected classes of proteins. We conclude that selection of appropriate negative controls is a critical aspect of proteomic experiments.

## Materials and Methods

### Strains

Worms were grown at 23°C on nematode growth medium (NGM) plates seeded with *Escherichia coli* OP50 as a food source. Imaging analysis was performed at the larval L4 stage. *C. elegans* strains used in this study can be found in Supplementary Table 1.

### Transgenic lines

Transgenic lines were prepared by gonadal microinjection of expression vectors for overexpression models or CRISPR/Cas9 for endogenous transgene expression or editing. Overexpression clones were made in the pSM vector^38^. Pan-neuronal overexpression was driven by the promoter rgef-1 and DA9-specific expression was driven by the mig-13 promoter. Standard techniques were used in the preparation of the plasmids and transgenic strains were prepared by microinjection using 1–5 ng/μl of plasmid DNA and coinjected with markers Podr-1::RFP at 100 ng/μl.

### Generation of neurexin TurboID-tagged by CRISPR/Cas9

Neurexin was TurboID-tagged by CRISPR-mediated insertion of TurboID into the endogenous neurexin genomic locus either just after the transmembrane domain (“post-TM”) or right before neurexin’s PDZ biding motif (“pre-PBM”) near the C terminus of the protein. To create the “pre-PBM” neurexin-TurboID strain used for the proteomics experiments in this study, the microinjection mix contained a crRNA with a guide RNA chosen close to the site of interest (3’ AAACGGAAACGGGAATGGG 5’), Alt-R® S.p. Cas9 Nuclease V3 (IDT, Cat. # 1081058) and a repair template generated by PCR that included the TurboID gene embedded with unc-119(+) cassette flanked by loxP sites within TurboID’s intron and a 96bp and 97bp homology arms to Cas9 cut site. DP38 [unc-119(ed3) III] strain was crossed with TV18675 (wyIs685 [Pmig-13::GFP::cla-1S + Pmig13::tdTomato::rab-3]) and the resulting strain PT23 [unc-119(ed3) III; nrx-1(kur2), wyIs685 V] was used for the injections. Transgenic animals were then selected based on behavioral rescue of the UNC phenotype by the expression of unc-119(+) and confirmed by PCR genotyping. Unc-119(+) cassette was then deleted by overexpression of Cre recombinase performed by microinjection of the plasmid pDD104 (Peft-3::Cre; Adgene). Genetic edited animals were selected based on Unc phenotype and confirmed again with PCR genotyping. Lastly, animals were out-crossed with N2 males to select away the unc-119(ed3) III allele resulting in the PTK31 [nrx-1(kur2), wyIs685 V] used for imaging and PTK57 [nrx-1(kur2) V] strain used for the proteomics in this study.

### Generation of neurexin-ΔPBM-TurboID by CRISPR/Cas9

Removal of the PDZ biding motif (PBM) from the neurexin-TurboID strain (PTK57) was performed by CRISPR-mediated deletion. For this purpose a co-CRISPR methodology^39^ was employed. PTK57 was injected with a mix containing crRNA targeting the PBM region (guide sequences used: 3’ TTTCTTCAATCAAAACTCAA 5’, 3’ AGAAAAAGGATTTTAAAGAG 5’ and 3’ GGTGGCACAGGAGGAACGGG 5’), a repair templated for the deletion with 100bp homology arms flanking the PBM, as well as a crRNA targeting the dpy-10 gene and its repair template^39^. Roller worms were then singled and genotyped for PBM deletion and these worms were subsequently passed to select away from the dpy-10 allele resulting in the PTK226 [nrx-1(kur43) V] strain used as a control in our proteomics experiments.

### Protein extraction for proteomics and western blotting

Protein extracts were prepared by harvesting synchronized worms enriched for adults with M9 onto a microcentrifuge tube followed by three M9 washes and two milli-q H_2_O washes. In the condition with added biotin, prior to the washes, worms were incubated at room temperature (22°C) in M9 buffer supplemented with 1 mM biotin, and *E. coli* OP50 for 2 h. After the washes, lysis buffer (150mM NaCl; 50mM Tris pH 8 and 0.1% NP-40) was added to the samples which were snap frozen in liquid nitrogen. This was followed by three cycles of pestle grinding/snap freezing and lastly by a 20,000g centrifugation at 4°C for 20 minutes. The protein content on the extracts was quantified using Pierce™ BCA Protein Assay Kit (Cat. #23225).

### Western Blotting

10μg of protein extracts were separated by 10% SDS-PAGE Tris-glycine polyacrylamide gel electrophoresis. 0.2 μm nitrocellulose membranes were used for the transfer in Towbin buffer for 4h at constant 280mA. Blots were incubated for 5 minutes with Ponceau S (0.1% (w/v) Ponceau S in 5% glacial acetic acid) for total protein visualization to control for possible loading differences. For immunodetection of biotinylated proteins, membranes were blocked in 7% milk in 1xTBS and 0.01% Tween-20 and streptavidin-HRP immunostaining (1:5000, Invitrogen cat. #19534-050) was performed at room temperature for 1h in blocking solution. After 3 washes with TBST, membranes were covered with SuperSignal™ West Femto Maximum Sensitive Substrate (Thermo Scientific, Cat. #34095) according to manufacturer’s instructions and chemiluminescence was then documented using Azure 600 Western Blot Imaging System (Azure Biosystems, Inc).

### Proteomics Streptavidin pull-downs and Mass Spectrometry

100 μg of protein extracts were incubated with freshly washed Pierce Streptavidin Plus Ultra-Link Resin (Thermo Scientific, Cat. #53117) in protein binding buffer [150 mM NaCl; 50 mM Tris pH 8; 10 μM ZnCl2; 0.5 mM DTT; 1:10 complete protease inhibitors (Sigma-Aldrich, Cat. #P2714); 10 mM sodium butyrate] for 6h at 4°C in a rotation wheel. Supernatant was discarded and streptavidin beads were resuspended in 100 μL of protein binding/wash buffer (350 mM NaCl; 50 mM Tris pH 8; 10 μM ZnCl2) followed by loading the samples into the desalting plate (Orochem OF1100 96-well plate) and 5 washes with protein binding/wash buffer. To reduce disulfide bonds, a 1h incubation at room temperature with 100 μL of 5 mM of DTT in 50 mM ammonium bicarbonate was done, which was followed by blocking reduced cysteine residues with 20 mM of iodoacetamide (100 μl/well) during 30 min in the dark also at room temperature. After blocking, flow-through was discarded and trypsin incubation (250 ng/well) was performed overnight with a 60% ACN in 0.1% TFA (25 μL) wash right after. Desalting was then performed as previously described^40^ followed by mass spectrometry. Briefly, samples were loaded onto a Dionex RSLC Ultimate 300 (Thermo Scientific), coupled online with an Orbitrap Fusion Lumos (Thermo Scientific). Chromatographic separation was performed with a two-column system, consisting of a C-18 trap cartridge (300 μm ID, 5 mm length) and a picofrit analytical column (75 μm ID, 25 cm length) packed in-house with reversed-phase Repro-Sil Pur C18-AQ 3 μm resin. Peptides were separated using a 60 min gradient from 4-30% buffer B (buffer A: 0.1% formic acid, buffer B: 80% acetonitrile + 0.1% formic acid) at a flow rate of 300 nl/min. The mass spectrometer was set to acquire spectra in a data-dependent acquisition (DDA) mode. Briefly, the full MS scan was set to 300-1200 m/z in the orbitrap with a resolution of 120,000 (at 200 m/z) and an AGC target of 5×10e5. MS/MS was performed in the ion trap using the top speed mode (2 secs), an AGC target of 1×10e4 and an HCD collision energy of 35. Raw files were searched using Proteome Discoverer software (v2.4, Thermo Scientific) using SEQUEST search engine and the SwissProt C. elegans database. The search for total proteome included variable modification of N-terminal acetylation, and fixed modification of carbamidomethyl cysteine. Trypsin was specified as the digestive enzyme with up to 2 missed cleavages allowed. Mass tolerance was set to 10 pm for precursor ions and 0.2 Da for product ions. Peptide and protein false discovery rate was set to 1%. Proteomics data transformation and normalization was performed as previously described^28^. Mass spectrometry raw files are deposited in the repository Chorus (https://chorusproject.org/) under the project number 1791.

### Immunohistochemistry

Immunohistochemistry was performed using the freeze-crack protocol described in www.wormbook.org with the following modifications. Ice cold 4% PFA was used as fixative solution with a 2h incubation at 4°C. This was followed by blocking with 1% Triton X-100, 1mM EDTA pH8, 0.1% BSA and 7% normal donkey serum in 1x PBS for 4h at room temperature. Incubation with mouse anti-BirA primary antibody (1:250, Abcam, Cat. #Ab232732) was performed overnight at 4°C. Secondary antibody incubation was also performed overnight at 4°C with donkey anti-mouse Alexa 647 (1:250, Invitrogen, Cat. # A-31571).

### Confocal microscopy

Imaging was performed at room temperature in live *C. elegans* grown at 23°C. An average of 20 mid-L4 stage hermaphrodite worms were paralyzed with 10 mM levamisole (Sigma-Aldrich) in M9 buffer, and mounted on 5% agar pads for imaging. Animal stage was determined based on the correct stage of vulval development using DIC optics. Images of fluorescently-tagged fusion proteins were captured with a Zeiss Axio Observer Z1 microscope with a Plan-Apochromat 63x or 40x 1.4NA objective and a Yokagawa spinning-disk unit attached to an EM-CCD camera.

### Image processing and data quantification

Using ImageJ (NIH), maximum-intensity projections were generated followed by cropping and straightening of the images. Puncta number was then quantified using a custom ROI-based MATLAB application (Image Processing Toolbox Release 2022a, The MathWorks, Inc., Natick, MA) using local mean thresholding and ROI watershed segmentation followed by parametric restriction to remove noise pixels. Image levels, whenever required, were adjusted in Adobe Photoshop to show relevant features. In such cases, any images being compared were treated in the same manner.

### Gene ontology (GO) enrichment analysis

GO analysis was performed using the enriched (upregulated) portion of the proteomics hits from the neurexin-TurboID strain samples when compared to the control samples. Figures in this manuscript focus on neurexin-TurboID vs. neurexin-ΔPBM-TurboID enriched hits as describet in the text. These hits were uploaded to https://wormbase.org/tools/enrichment/tea/tea.cgi server form GO enrichment analysis.

### Statistical analyses

GraphPad Prism 9.0 (GraphPad Software, La Jolla, CA, USA) software was used for the statistical analyses. Student’s t-test was used to test for significance compared to controls and all data are represented as mean ± SEM, and significance is defined as *p < 0.05, **p < 0.01, or ***p < 0.001, unless otherwise noted.

Volcano and Semi-Volcano plots were constructed using two-sample t-test to evaluate differential protein levels between conditions followed by plotting the p-value (-log10) against the fold change (log2) (MATLAB and Bioinformatics Toolbox Release 2022a, The MathWorks, Inc., Natick, MA).

## Acknowledgements

We thank Dr. XiaoGuang Wang for generating the transgenic rescue strains and “post-TM” TurboID crispr strain. Some strains were provided by the CGC, which is funded by NIH Office of Research Infrastructure Programs (P40 OD010440), Additional strains were provided by the National BioResource Project (Japan). PTK and MSP were funded by the Simons Foundation (SFARI pilot award) and the Mathers Foundation. SS gratefully acknowledges for financial support AFAR (Sagol Network GerOmics award), Deerfield (Xseed award), Relay Therapeutics, Merck and the Einstein-Mount Sinai Diabetes Research Center.

## Figure Legends

**Supplementary Figure 1.**
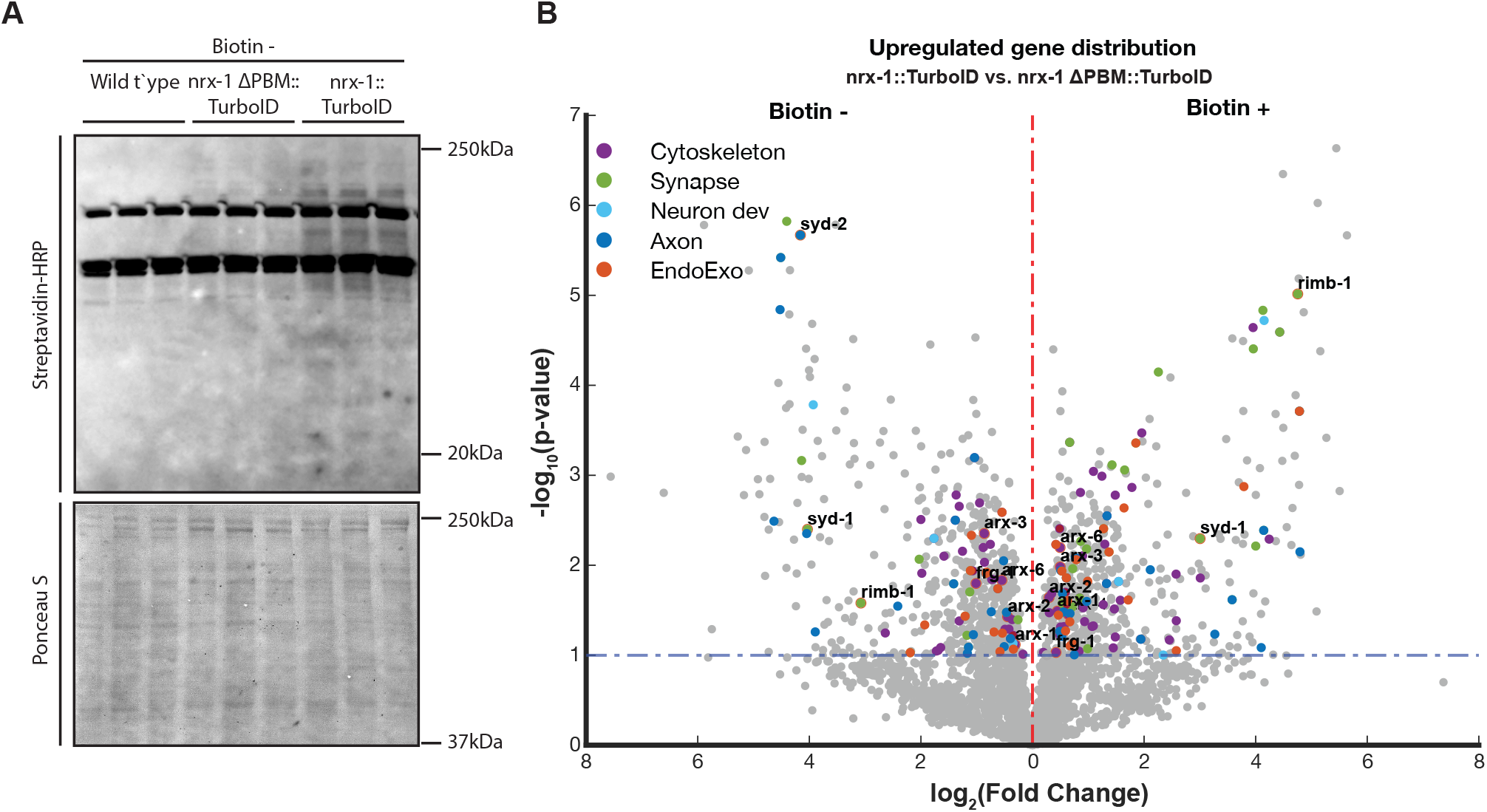
Assessment of control strains. **A**. Western blot of biotin-tagged proteins of our experimental strain (neurexin-TurboID, right columns) compared to two controls (wild type, left columns and neurexin-ΔPBM-TurboID, middle columns) in the basal, non-enriched Biotin condition. Total protein levels (as assessed by Ponceau staining, lower blot) were used as a loading control. **B**. Volcano plot of genes corresponding to the proteins enriched in our experimental strain (neurexin-TurboID) compared to control (neurexin-ΔPBM-TurboID), in either the basal (left) or enriched (right) biotin conditions.

**Supplementary Figure 2.**
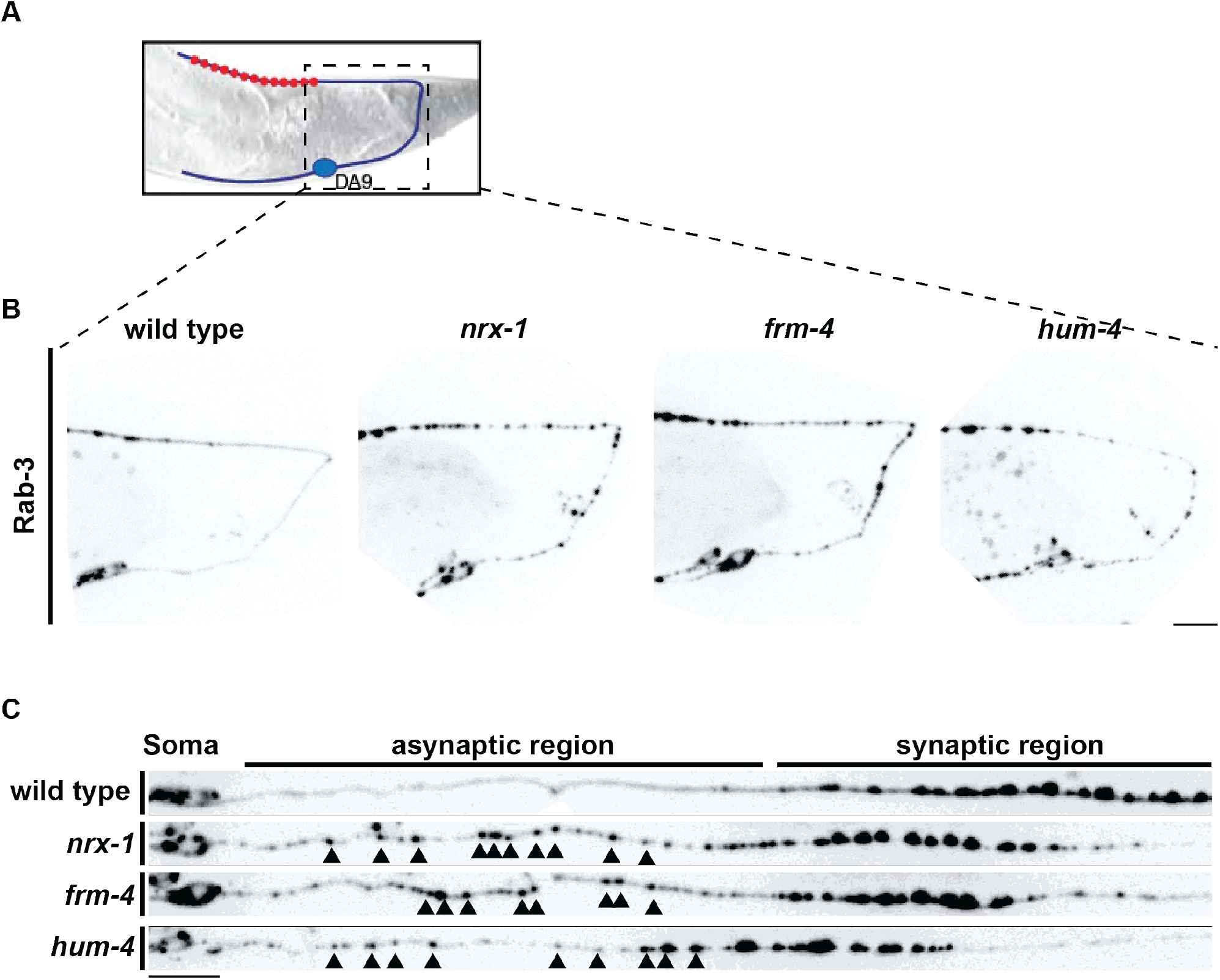
Neurexin RAB-3 phenotype present in some candidate interactors. **A**. Schematic of the worm tail showing the region of the images. **B**. Images of the DA9 motor neuron showing RAB-3-TdTomato fluorescence, which is normally restricted to the synaptic region in wild type (left) but reveals small asynaptic puncta in neurexin mutants (middle) and frm-4 mutants (right). **C**. Cropped and straightened images of the axon, starting at the cell body and ending at the synaptic domain. Arrowheads display examples of asynaptic RAB-3 puncta not present in wild type. Scale bars: 10 μm.

**Supplementary Figure 3.**
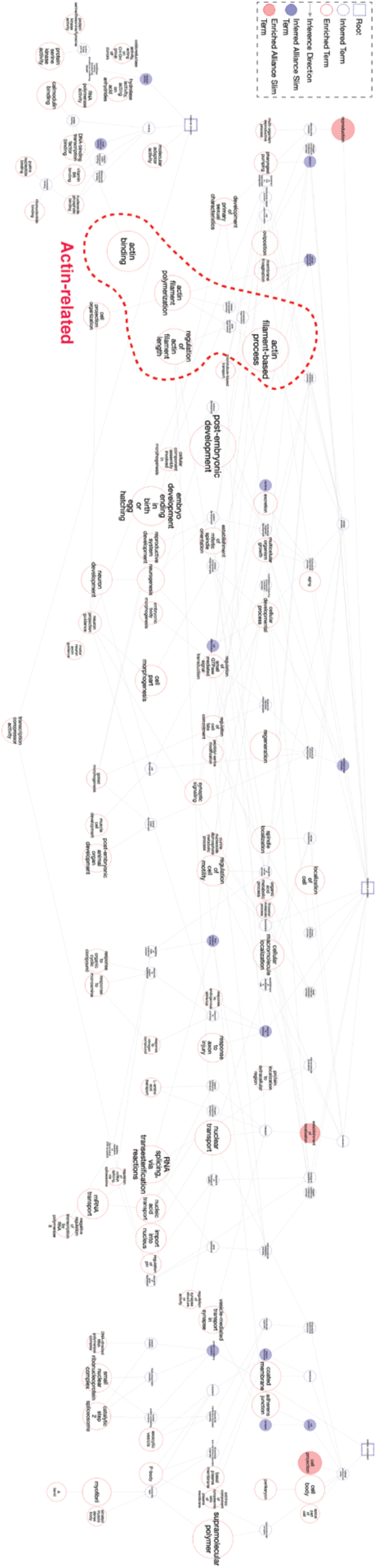
GO analysis connectome showing prominent enrichment of actin-related proteins. Connectome displaying the different GO terms found to be enriched in the samples. Actin-related terms are highlighted by the dotted red segment ROI of the map.

## Notes

### Competing Interest Statement

The authors have declared no competing interest.

### Summary of Updates

Additional experimental data on neurexin's link to actin polymerization (new figure 5) and information about how to access the raw mass spec data.

